# Ancestral state reconstruction suggests repeated losses of recruitment communication during ant evolution (Hymenoptera: Formicidae)

**DOI:** 10.1101/2022.05.18.492496

**Authors:** Simone M Glaser, Christoph Grüter

## Abstract

Eusocial insects have evolved different strategies to share information about their environment and workers can recruit nestmates to food sources or new nest sites. Ants are the most species-rich social insect group and are known to use pheromones, visual and tactile signals to communicate and inform nestmates about resources. However, how these different strategies evolved and whether there was a predominant evolutionary sequence that led to present day recruitment strategies is not well understood. In our study we explored two competing hypotheses about the ancestral recruitment communication: (1) ant ancestors did not recruit nestmates and species evolved more complex recruitment strategies over time vs. (2) early ants used mass-recruitment, which was lost repeatedly in some lineages. We combined an extensive search of the scientific literature and ancestral state reconstruction to estimate the ancestral recruitment strategy, focusing on the categories (i) no recruitment, (ii) tandem running, (iii) group-recruitment and (iv) chemical mass-recruitment. Stochastic character mapping suggests that mass-recruitment was ancestral in ants (59-61%), whereas “no recruitment” was unlikely to be the ancestral condition (21%). Similarly, marginal ancestral state reconstruction suggests that mass-recruitment (44-81%) or group-recruitment (48-50%) represented the original state. Our results are consistent with the finding that early ants lived in colonies containing up to several thousand individuals, which are typically associated with mass-recruiting in ants. However, our ability to robustly identify patterns in the evolution of communication in ants remains hampered by a lack of natural history information for most ant species.

## Introduction

Communication about resources is widespread in social insects and different species have evolved a variety of strategies to communicate with other individuals (Czaczkes et al., 2015; Franklin, 2014; Grüter & Leadbeater, 2014; Hölldobler & Wilson, 1990; Leadbeater & Chittka, 2007; von Frisch, 1967; Wilson, 1971; Grüter, 2020). Senders can use visual, tactile (body contact in honeybees, Grooters, 1987; antennation, Hölldobler & Wilson, 1990; von Frisch, 1967) or chemical (e.g. pheromones, Czaczkes et al., 2015; Hölldobler & Wilson, 1990) signals to share information about food locations, nesting locations, dangers or the needs of individuals and the colony.

Ants (Family: Formicidae) are an extraordinarily diverse, widespread and ecologically important group of social insects, containing over 13,000 extant species (Reeves & Moreau, 2019). Ants probably first appeared in the early Cretaceous, more than 100 million years ago, most likely from a lineage of wasps (Hölldobler & Wilson 2009; Moreau et al., 2006; Ward, 2014). Ants have evolved a variety of different communication strategies, some of which are behaviourally highly complex (Hölldobler, 1999). There are also species that employ several strategies during recruitment (see below).

Several classification systems for recruitment behaviours have been proposed (Beckers et al., 1989; Jaffe, 1984; Lanan, 2014). The most common strategies are the use of pheromone trails (Czaczkes et al., 2015; Hölldobler & Wilson, 1990, 2009), group-recruitment (Hölldobler, 1971; Liefke et al., 2001), tandem recruitment (Franklin, 2014; Glaser & Grüter, 2018; Grüter et al., 2018) or no recruitment at all (solitary/individual) (Beckers et al., 1989; Jaffe, 1984). In tandem recruitment, a scout with knowledge of a valuable resource recruits a single nestmate to that resource. The follower maintains contact with the leader by antennating the gaster and hind legs of the leader. In group-recruitment, a scout leads a group of up to thirty nestmates to a resource (Beckers et al. 1989; Liefke et al., 2001). A short-lived, volatile pheromone emitted by the leader helps recruited ants stay close to the leader (Möglich et al., 1974). In mass-recruitment, successful scouts lay pheromone trails between the nest and the resource, which is followed and strengthened by nestmates. Some species use a particular strategy in one context but not in another. For example, *Neoponera* or *Diacamma* species perform tandem runs when they relocate to a new nest site, but when they are foraging for prey they do so solitarily (Fresneau, 1985; Kaur et al., 2017). Furthermore, recruitment communication can depend on the type of food source that is collected. When collecting small prey, ants often do not recruit nestmates as they can carry home the food item by themselves, but use recruitment communication when finding larger resources, like honey dew-secreting aphid colonies or large prey items (Czaczkes et al., 2011; Czaczkes & Ratnieks, 2012; Detrain & Deneubourg, 2008; Lach, 2005). These examples highlight that the use of recruitment communication often depends on the ecological context.

Species with small colonies (<1000 individuals) often do not use recruitment or they perform tandem runs, whereas medium size colonies (up to a few thousand individuals) tend to recruit nestmates by group-recruitment or by pheromone trails and large colonies use mainly pheromone trails (Beckers et al., 1989). Recruitment *via* pheromone trails requires colonies to have a minimum number of foragers to deposit pheromones in order to maintain the trail (Beekman et al., 2001). The link between colony size and recruitment strategy is not rigid, however, and species with similar colony sizes can differ in the strategies they employ (Beckers et al., 1989).

This diversity in recruitment strategies raises the question how these strategies evolved and whether certain forms of recruitment tend to precede other methods. One hypothesis is that recruitment communication increased in complexity over evolutionary time. According to this scenario, early ants would have foraged solitarily, like many present-day ponerine ants (Beckers et al., 1989; Maschwitz & Schönegge, 1983; Villet, 1990). Subsequently, small-scale communication mechanisms evolved, like tandem-running and group-recruitment. From these forms of communication, mass-recruitment with longer-lasting chemical trails may have evolved (Beckers et al., 1989; Hölldobler & Wilson, 1990; Traniello, 1989). Hingston (1929) and Wilson (1959), for instance, suggested that tandem running is an ancestral form of communication that was used by early ants. Short-lived pheromones released by the tandem leader play a potentially important role for the cohesion between the leader and follower in a tandem run (Basari et al., 2014). It would then have been a small evolutionary step to produce longer lasting trail pheromones that allowed mass-recruitment in species with larger colony sizes. The hypothesis of an increase in scale – from small-scale to large-scale recruitment – has recently been supported by Reeves & Moreau (2019) who suggest solitary foraging as the ancestral recruitment strategy.

A phylogenetic analysis by Burchill & Moreau (2016), on the other hand, suggested that early ant species had medium colony sizes, with colonies containing up to several thousand individuals, which is typically associated with mass-recruitment in extant ants (Beckers et al., 1989). This suggests that mass-recruitment may have been a more likely strategy used by early ants. Following this argument, recruitment would have been lost over time in some lineages as ant species with small colony sizes evolved (Burchill & Moreau 2016). The antiquity of tandem running has also been questioned by the fact that this behaviour is found in species that are considered more derived, such as in *Temnothorax* and *Leptothorax* (Planque et al. 2010). This suggests that tandem running may be a derived behaviour, at least in some groups.

The main aim of our study was to estimate the ancestral state of recruitment communication of the Formicidae. While Reeves & Moreau (2019) focused on recruitment communication during foraging, we also considered whether species use recruitment during emigrations. We included these cases because numerous ant species do not communicate during foraging but use recruitment communication in other ecological situations (Fresneau, 1985; Grüter et al., 2018; Kaur et al., 2017). The value of communication in foraging depends on the foraging ecology of a species, such as the kind of food that is exploited and food source distribution (Anna Dornhaus et al., 2006; I’Anson Price et al., 2019; Sherman & Visscher, 2002). Thus, the strategies used during foraging reflect foraging ecology and provide an incomplete picture of the recruitment strategies used by a species. To better understand the evolution of recruitment communication mechanisms, it is instructive to consider whether a species uses recruitment communication, irrespective of the type of resource that is exploited. In addition, we explored whether tandem running was indeed an early recruitment strategy that preceded group- and mass-recruitment (Hingston, 1929; Wilson, 1959).

## Material & Methods

### Literature research for recruitment strategies

Data were collected on the recruitment strategies used by extant ant species *via* an extensive search of the published scientific literature (from October 2019 to March 2020). For many ant species, information about recruitment was collected from reviews or articles about recruitment (Beckers et al., 1989; Jaffe, 1984; Silvestre et al., 1999). Furthermore, we searched in Google Scholar using the search terms (ant species or genus in combination with “recruit”, “forag”, “prey”, “individual”, “solitary”, “tandem”, “group”, “trail”, “pheromone”). We included species-level information when the recruitment strategy was described based on observations or collected in controlled experiments. Data were coded as discrete character traits. Each species was allocated to one of four different recruitment strategies, similar to Jaffé (1984), Beckers et al. (1989) and Lanan (2014): no recruitment, tandem running, group-recruitment and mass-recruitment (Table 1). Reeves & Moreau (2019) found that the different classification systems led to very similar outcomes in their ancestral state reconstruction.

**Table 1.** Recruitment classifications and definitions, largely based on Jaffé (1984).

| Recruitment strategy | Definition |
| --- | --- |
| **Solitary/individual** | No recruitment, no information transfer between nestmates |
| **Tandem running** | A single ant (scout) attracts a single nestmate using antennal contact and then physically leads a nestmate to the goal. Physical contact is often maintained between scout and nestmate, chemical signals may be used |
| **Group-recruitment** | A scout recruits “up to thirty nestmates” and leads them to the goal. Chemical signals are often used for short-distance attraction but physical contact between scout and the group is also used |
| **Mass-recruitment** | Groups are guided via chemical trails alone. Large numbers of ants can be recruited by a small number of recruiters. Chemical trails are laid on the substrate. |

The literature search highlighted that there is a relative scarcity of detailed information about foraging strategies in the ant literature (see also Reeves & Moreau 2019). There are numerous examples of studies that mention recruitment of nestmates, but without providing details of the strategy and the context when this was observed. These studies were not included in our analysis. If possible, we also collected and compared the recruitment data to Reeves and Moreau (2019), who collected data on ant foraging strategies. In some cases, we were unable to recover the information from the cited primary literature (e.g. *Buniapone amblyops, Mayaponera constricta* or *Megaponera analis*). *Tetramorium caespitum* uses both group and mass-recruitment (Collignon & Detrain, 2010). We considered this to be a mass-recruiting species for our analysis. Additionally, we used a subset that included only ant species that perform tandem runs and analysed if recruitment to nest sites or food sources was more likely to be the ancestral state.

### Phylogenetic comparative methods

We modified the phylogenetic tree of Branstetter et al. (2017), which contains ∼1000 ant species and is a phylogram based on molecular data. In our literature research we found information for 161 species, 82 genera and 11 sub-families (Table S1) that were also present in the phylogenetic tree. Overall, the species included in our study represent 25% of genera and 65% of sub-families. Species with no recruitment data available were removed from the dataset with the *drop*.*tip* function in the R package “ape” (Paradis & Schliep, 2019).

We performed marginal ancestral state reconstructions (ASR) for the dataset, using the functions *fitMk* and *ace* from the R package “phytools” (Revell, 2012) and “ape” (Paradis & Schliep, 2019) to estimate the transition rates and the ancestral states for our tested character using a maximum likelihood (ML) approach. The *fitMk* function assumes that the probability to change from one state to another depends only on the current state and not on the state that has come before. Furthermore, every character state is equally likely to change to one of the other states. The *ace* function utilizes marginal reconstruction and returns the marginal ancestral state likelihood of all nodes within a phylogeny.

Additionally, we performed a stochastic character mapping (SCM) by using *make*.*simmap* from the R package “phytools”. For the stochastic reconstructions of character states we used an MCMC approach, to explore the posterior probabilities of all nodes and provided the number of changes between the character states (1000 simulations performed).

Three commonly used transition rate models were analysed for the ancestral state reconstructions: “equal rates” (ER), “symmetrical rates” (SYM) and “all rates different” (ARD) with names referring to transition rates between each state. We used the Akaike information criterion (AIC) values corrected for small sample sizes (AICc values) for the three transition rates. We calculated the AIC-weights which standardize the AIC scores of the fitted models and measured the relative weight of evidence for the three models used in our data (Harmon, 2019). We visualized the results by mapping the ancestral state on the phylogeny with the function *plotTree*.

## Results

### Evolution of recruitment strategy

The ancestral state reconstruction results were mapped to our phylogeny (Fig. 1). The log-likelihood values, AIC values, AICc values and the number of free parameters per model are presented in Table 2. We compared the AIC and AICc values, which revealed that simple ER models were inferior and, thus, were rejected. A “symmetric model” and “all-rates-different model” best explained our transition between recruitment states. Thus, both transition models were used to analyse the ancestral state of recruitment strategies in ants.

**Table 2.** Results of the transition rate models. Log likelihoods, Akaike information criterion values, number of free parameters and Akaike-weights are shown.

| Model | LogL | AIC | AICc | free parameters | AICcW |
| --- | --- | --- | --- | --- | --- |
| ER | -170.0 | 336.9 | 342.0 | 1 | 0.03 |
| SYM | -161.4 | 331.6 | 335.4 | 6 | 0.45 |
| ARD | -155.7 | 331.4 | 337.6 | 12 | 0.52 |

**Figure 1.**
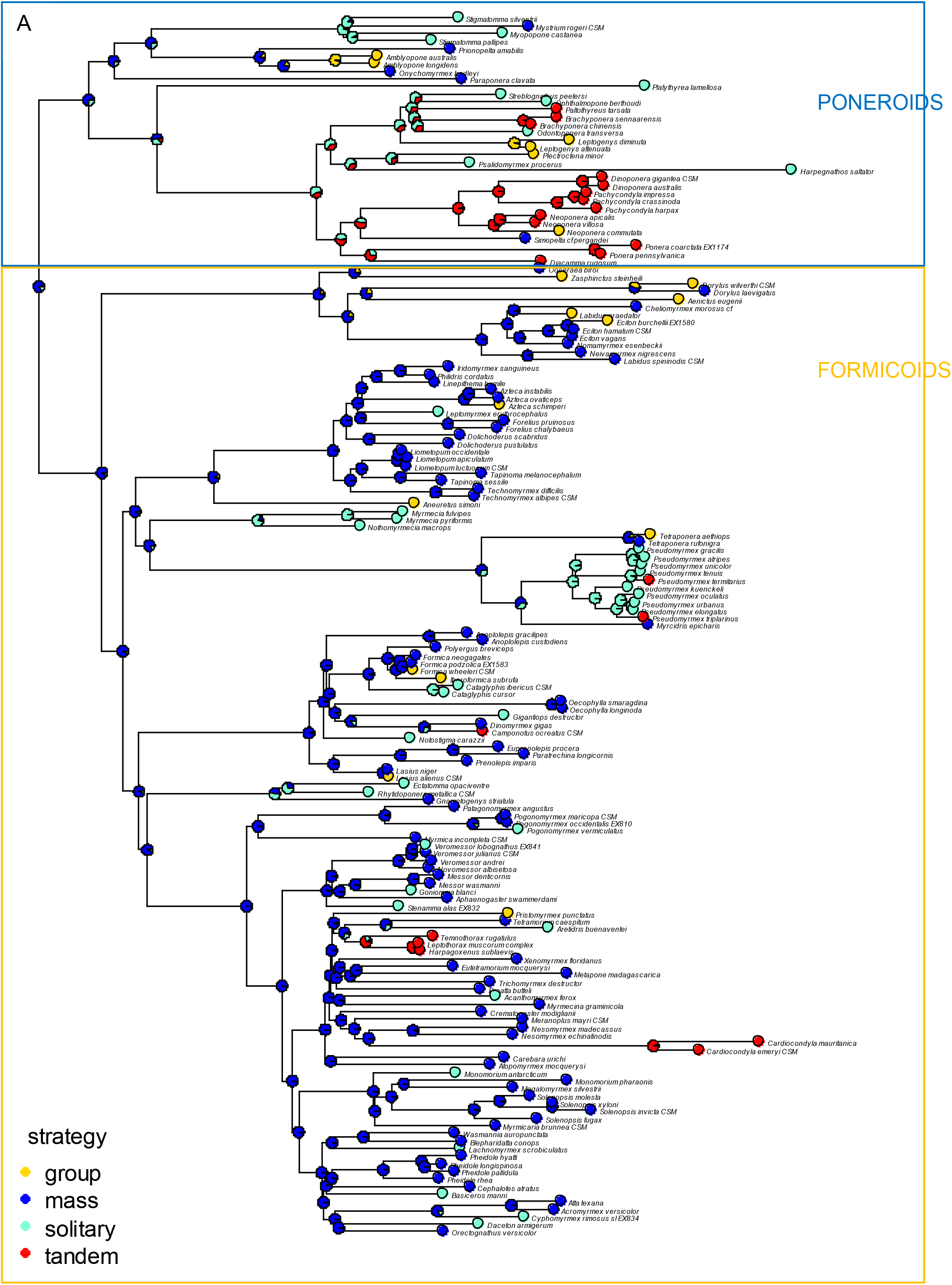
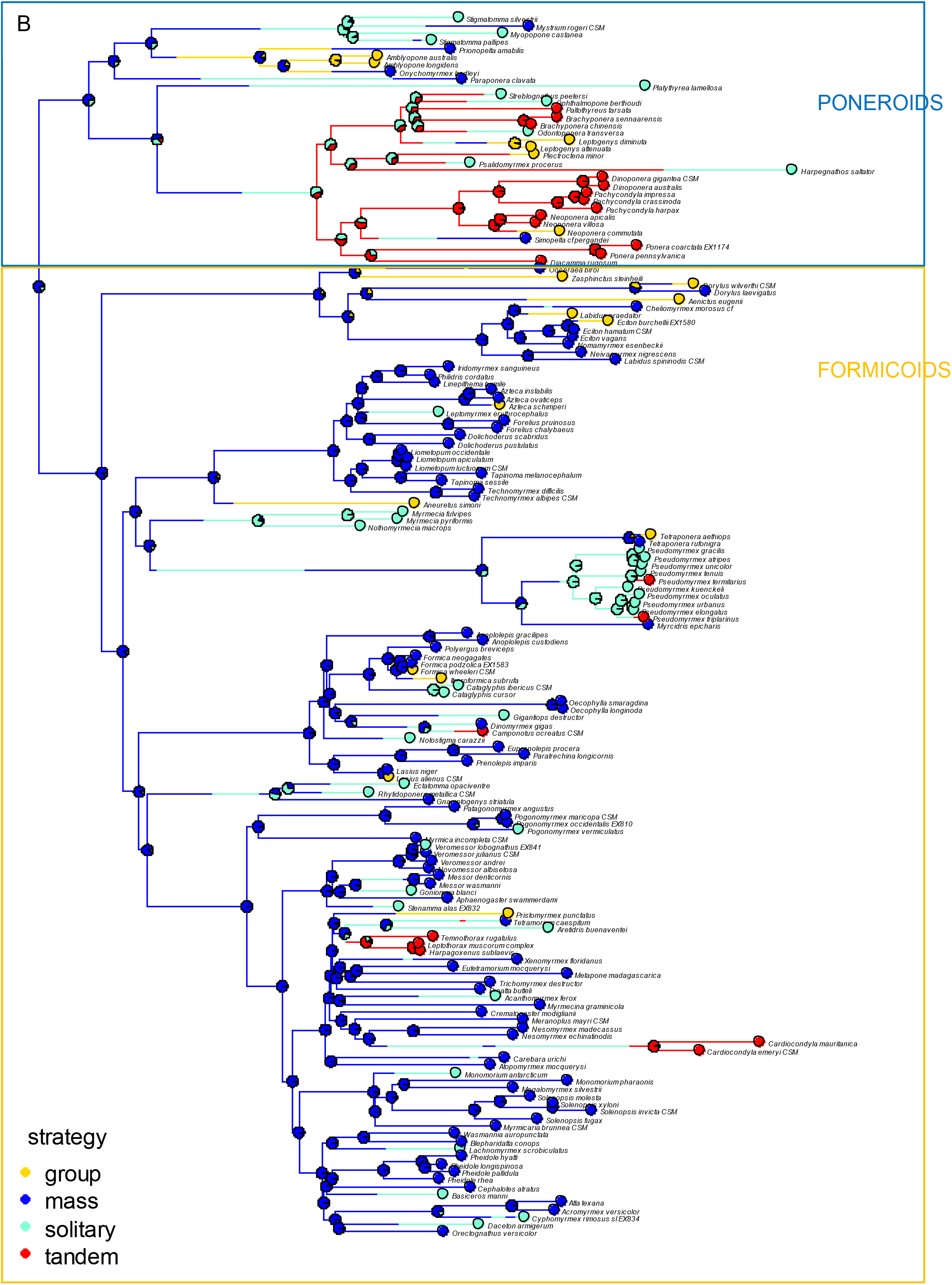
Ant phylogeny including recruitment strategy and (**A**) marginal ancestral state reconstruction or (**B**) stochastic character mapping. Nodes provide estimates based on Markov chain models. The phylogeny is based on Branstetter et al. (2017).

Marginal ancestral state reconstruction analyses and stochastic character mapping both suggest that mass-recruitment is the most probable strategy at the root of the phylogeny (71.2% and 60.2%, respectively) (Table 3). Also, internal nodes (lineage splitting events) were dominated by high probabilities for the mass-recruitment category. Mass-recruitment was the most likely ancestral state in both the Poneroids and the Formicoids (Figure 1). The stochastic character mapping revealed that there were an estimated 81.3 changes between recruitment strategies (Table 4). The most common transitions were from mass-recruitment to solitary/individual behaviour (33.7%) or to group-recruitment (18.5%) (Figure 2). Furthermore, there were transitions from no recruitment to tandem running (14.8%) or mass-recruitment (12.7%). Tandem running evolved several times independently in the subfamilies Ponerinae, Pseudomyrmecinae, Formicinae and Myrmecinae. Furthermore, it seems that recruitment was lost at least once in all subfamilies, except in the Dorylinae (army ants). Similarly, we found group-recruitment in nearly all included sub-families, except in the Paraponerinae. The Myrmeciinae were the only group without species that perform mass-recruitment.

**Table 3.**
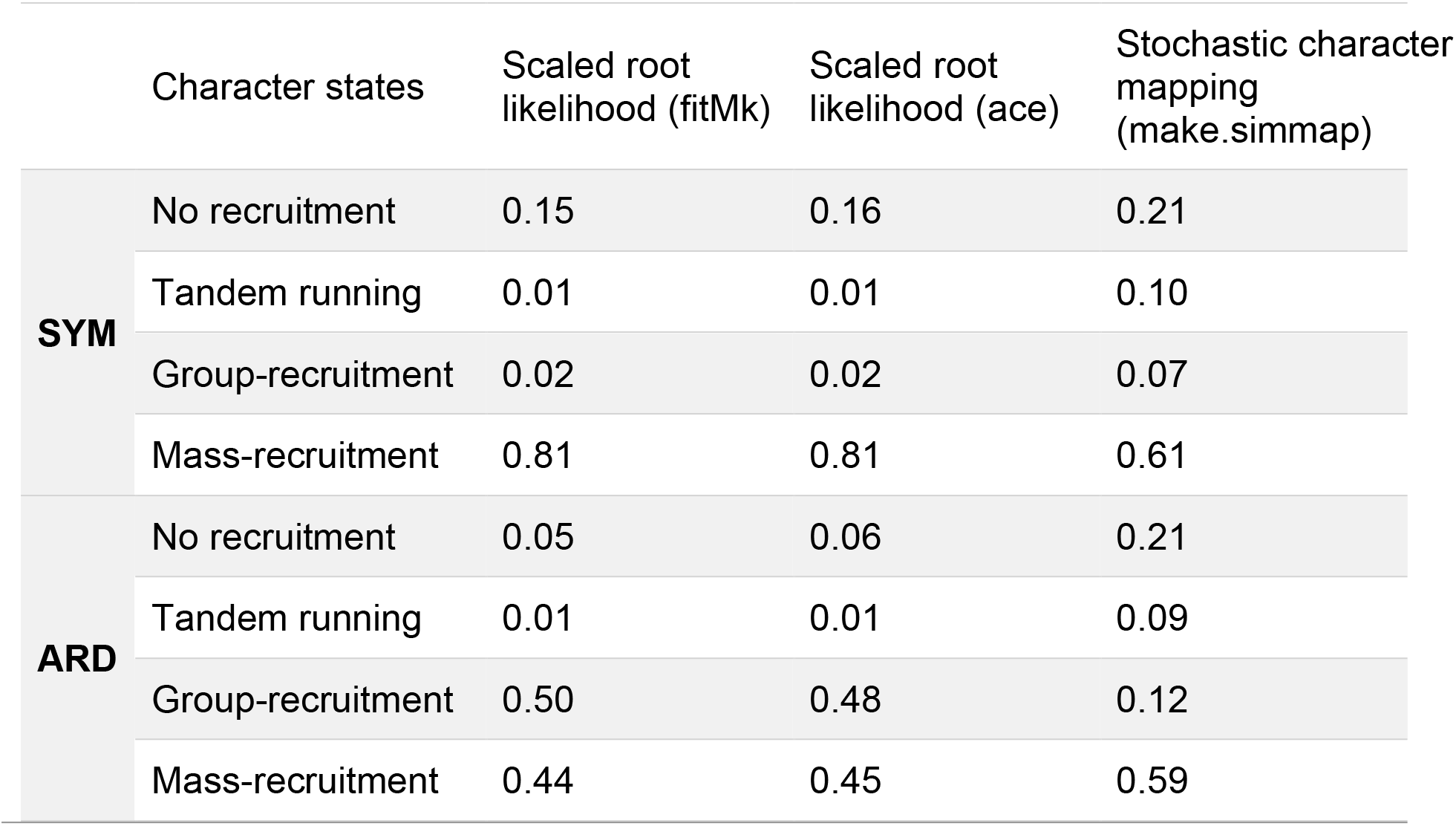
Ancestral character estimation using marginal ancestral state reconstruction and stochastic character mapping. Values represent likelihoods of recruitment strategies at the root.

**Table 4.** Changes from stochastic character mapping. GR = group-recruitment, MS = mass-recruitment, NR = no recruitment, TR = tandem running

|  | **SYM** |  | **ARD** |  |
| --- | --- | --- | --- | --- |
| **Total changes** | 75.783 |  | 104.472 |  |
| **Type** | Number | Percentage | Number | Percentage |
| **GR → MS** | 3.44 | 4.5% | 32.013 | 30.6% |
| **GR → NR** | 0.595 | 0.7% | 0 | 0% |
| **GR → TR** | 1.224 | 1.6% | 0 | 0% |
| **MS → GR** | 14.902 | 19.7% | 23.572 | 22.6% |
| **MS → NR** | 26.57 | 35.1% | 27.514 | 26.3% |
| **MS → TR** | 0.489 | 0.6% | 0.952 | 0.9% |
| **NR → GR** | 1.412 | 1.9% | 7.409 | 7.1% |
| **NR → MS** | 10.061 | 13.3% | 0 | 0% |
| **NR → TR** | 9.8 | 12.9% | 9.572 | 9.2% |
| TR → GR | 2.626 | 3.4% | 3.44 | 3.3% |
| TR → MS | 0.117 | 0.2% | 0 | 0% |
| TR → NR | 4.547 | 6.0% | 0 | 0% |

**Figure 2.**
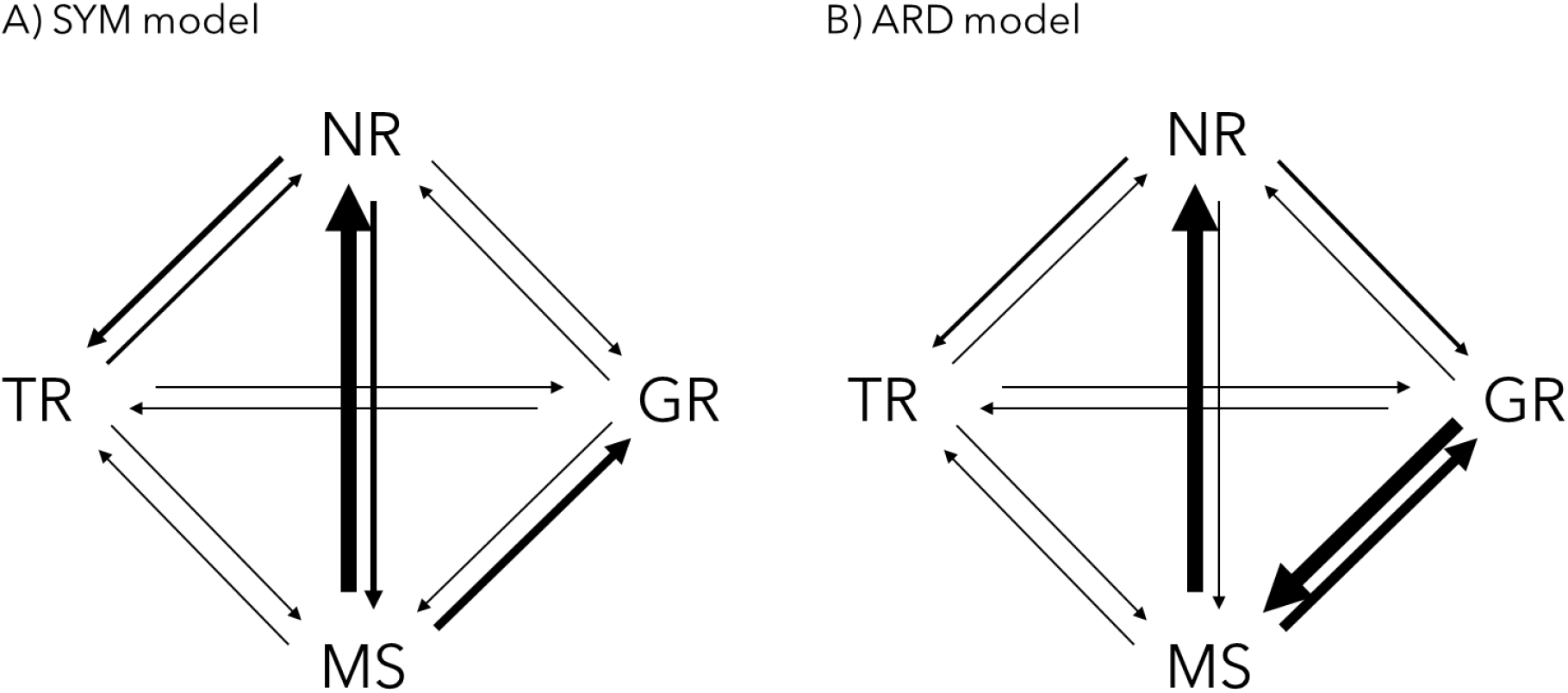
Transitions between states from stochastic character mapping for the methods SYM and ARD. The thickness of the arrow reflects the relative commonness of a transition. GR = group-recruitment, MS = mass-recruitment, NR = no recruitment, TR = tandem running

### Evolution of tandem running

Twenty-one species included in our study perform tandem runs. For some species it is not known if they recruit to food sources and during nest relocations. For other species it is known that they perform tandem runs to new nest sites, but forage solitarily for food sources (*Neoponera, Diacamma* or *Paltothyreus*) (Table S1).

After mapping the recruitment strategy onto the phylogenetic tree (Figure 3), we found that all species that can perform tandem runs do so during colony emigrations to new nest sites. Several species (43%) that use recruitment via tandem running do not do so during foraging.

**Figure 3.**
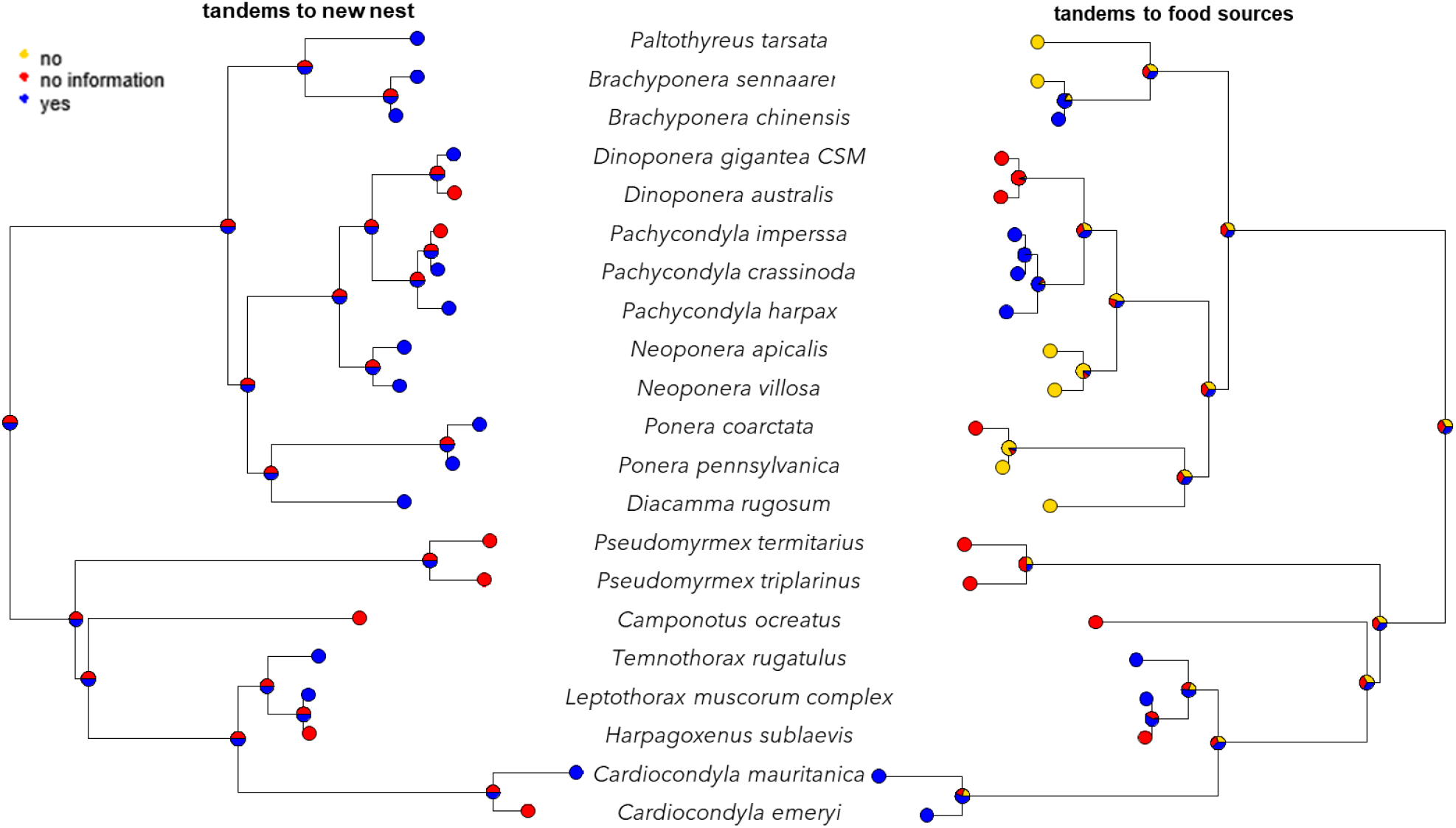
Ancestral state reconstruction for ant species that perform tandem running to either new nest sites, food sources or both. Nodes provide estimates based on Markov chain models.

## Discussion

Our ancestral state analyses indicate that mass-recruitment or group-recruitment were likely used for recruitment by the last common ancestor of present-day ants. During the course of their evolutionary history, all included subfamilies show switches from mass-recruitment to other recruitment strategies. Strikingly, most transitions occurred from mass recruitment to group or no recruitment. The repeated loss of communication seems puzzling, but probably coincided with the emergence of lineages with small colony sizes (e.g. in Amblyoponinae, Myrmeciinae, Ponerinae, Pseudomyrmecinae) (Burchill & Moreau 2016), where communication is less beneficial (Beckers et al. 1989). Transitions between mass- and group-recruitment were also more frequent than between tandem running and no recruitment. Tandem running evolved independently in 4 of 11 subfamilies. Furthermore, there were several transitions from no recruitment to tandem running or mass-recruitment. These findings highlight that recruitment strategies are an evolutionarily flexible and reversible trait.

Many lineages appear to have lost the ability to recruit nestmates. The putative loss of chemical mass-recruitment seems puzzling at first, but could be explained by the constraints of living in a small colony (Beckers et al., 1989; Dornhaus et al., 2012) and/or a switch to a diet or foraging strategy that does not require pheromone trails. Smaller colonies tend to exploit resources solitarily or they use tandem running. One reason could be that smaller colonies do not have sufficiently large colonies to maintain pheromone trails (Beckers et al., 1989; Beekman et al., 2001; Planqué et al., 2010). A recent study has suggested that the ancestral Formicidae had medium colony sizes containing up to several thousand individuals (Burchill & Moreau, 2016). These findings in combination with the findings that medium sized colonies often use group or mass-recruitment (Beckers et al. 1989) are consistent with our results that the best-supported ancestral recruitment strategy was mass-or group-recruitment.

Another reason for the loss of mass-recruitment could be that recruitment by pheromone trails can have disadvantages in changing foraging landscapes. Pheromone trails can persist for relatively long periods (up to several hours), which can make it difficult for the colony to re-allocate workers to a newly available higher-quality resource due to the strong positive feedback created by a pheromone trail (Beckers et al., 1989; Grüter et al. 2012; Czaczkes et al., 2015; I’Anson Price et al., 2016). This makes recruitment less flexible and colonies are more likely to miss out on new food sources when the environment changes.

Reeves & Moreau (2019) found evidence that solitary foraging, rather than mass recruitment, represented the original state in terms of recruitment strategies. Our and their results are not necessarily contradictory. In our study, we considered whether a species uses recruitment communication during colony emigrations and/or during foraging. It is well known that foraging strategies in ants strongly depend on the foraging ecology (Davidson, 1977; Dejean et al., 2012; Dornhaus et al., 2006; Hölldobler & Wilson, 1990; Lanan, 2014). For example, in many ant species, foragers follow a solitary foraging strategy when collecting insect prey, but they use recruitment communication when the colony emigrates to a new nest-site (Lanan, 2014). In other words, these species possess the ability to recruit, but foragers do not perform recruitment because this would not be an adaptive strategy given their foraging ecology.

The observation that numerous species recruit to new nest sites, but do not use recruitment communication during foraging (e.g. *Neoponera* or *Diacamma* species, Fresneau, 1985; Grüter et al., 2018) raises the question if recruitment communication evolved first to help colonies during emigrations rather than to communicate the location of food sources, as has also been suggested in the case of the honeybee waggle dance (Beekman et al. 2008; I’Anson Price & Grüter, 2015). This seems plausible given that during nest relocations of cavity nesting species, nest locations have to be communicated very precisely. If the old nest is damaged or destroyed, a fast and precise relocation is critical (Dornhaus et al., 2004; Franks et al., 2003). During foraging, on the other hand, communication might often be less important. Especially when food sources are abundant and evenly distributed, communication might not provide benefits or even be disadvantageous due to time costs (Dechaume-Moncharmont et al., 2005; Dornhaus et al., 2006; Goy et al., 2021; I’Anson Price et al., 2019). The hypothesis that recruitment evolved first in colony emigrations and was co-opted by some species in a foraging context is also supported by our results suggesting group-or mass-recruitment as the ancestral state and those of Reeves & Moreau (2019) who suggested solitary foraging as the ancestral condition.

It has been suggested that tandem running is a “primitive” recruitment strategy (*i*.*e*. ancestral) (Hingston 1929; Hölldobler et al., 1974; Schultheiss et al., 2015; Wilson, 1959). Our results do not support this view. We found that tandem running evolved repeatedly and independently in the subfamilies Ponerinae, Pseudomyrmecinae, Formicinae and Myrmecinae. Transitions to tandem running occurred most often from no recruitment and, more rarely, from mass or group-recruitment. One benefit of tandem running is that it allows small colonies to defend resources against competitors when competition for nest sites or food sources is intense(Glaser et al., 2021).

In summary, our results suggest that mass or group-recruitment were the most likely recruitment strategies used by the last common ancestor of present-day ants. There were repeated, independent transitions to different strategies, such as tandem running or no recruitment, but also transitions back to group or mass-recruitment. It should be noted that our analysis is restricted to a small proportion of ant species and we currently lack information about recruitment behaviours for the majority of species (see also Reeves & Moreau 2019). We echo the call of Reeves & Moreau (2019) to pay more attention to ant behaviour and ecology as this will allow us to better understand the links between different traits of ant behaviour, ecology and natural history.

## Acknowledgement

We thank the Department of Behavioural Ecology and Social Evolution, Johannes Gutenberg-University Mainz for feedback and advice. C.G. and S.M.G. were funded by the German Research Foundation (DFG: GR 4986/1-1).

